# Inhibition of striatal indirect pathway during second postnatal week leads to long lasting deficits in motivated behavior

**DOI:** 10.1101/2024.03.29.586960

**Authors:** Pedro R. Olivetti, Arturo Torres-Herraez, Ricardo Raudales, Mary-Elena Sumerau, Sinead Moyles, Peter Balsam, Christoph Kellendonk

## Abstract

Schizophrenia is a neuropsychiatric disorder with postulated neurodevelopmental etiology. Genetic and imaging studies have shown enhanced dopamine and D2 receptor occupancy in the striatum of patients with schizophrenia. However, whether alterations in postnatal striatal dopamine can lead to long lasting changes in brain function and behavior is still unclear. Here, we approximated striatal D2R hyperfunction in mice via designer receptor mediated activation of inhibitory Gi-protein signaling during a defined postnatal time window. We found that Gi-mediated inhibition of the indirect pathway during postnatal day 8-15 led to long lasting decreases in locomotor activity and motivated behavior measured in the adult animal. In vivo photometry further showed that the motivational deficit was associated with an attenuated adaptation of cue-evoked dopamine levels to changes in effort requirements. These data establish a sensitive time window of D2R-regulated striatal development with long lasting impacts on neuronal function and behavior.

## Introduction

Several neuropsychiatric disorders including schizophrenia (SCZ) are thought to have a neurodevelopmental origin but the understanding of neurodevelopment in the context of disease is still limited. Affecting multiple dimensions of neural function, SCZ is broadly defined by the emergence, starting in late adolescence, of hallucinations, delusions, and disorganized thinking (positive symptoms); deficits of working memory and cognitive flexibility; as well as motivation, sociability, and self-care (negative symptoms). Positive symptoms are treated primarily by pharmacological antagonism of striatal dopamine D2 receptors (D2R) and approximately 60% of individuals show a response to treatment. However, negative symptoms, especially motivation, generally do not respond well to antipsychotic medication, often leaving chronic patients with a long-term residual disease burden that greatly limits their ability to engage in their communities and participate in educational or professional activities [1].

Bridging this gap hinges on deepening our understanding of molecular and neurocircuit mechanisms underlying the regulation of motivated behavior in SCZ. Genetic and environmental studies suggest a neurodevelopmental origin of schizophrenia since many schizophrenia associated genes affect brain developments and external risk factors have been shown to act during embryonic and postnatal development [2–5].

At the pathophysiological level, alterations in dopamine (DA) neurotransmission and signaling through dopamine D2 receptors (D2Rs) remain key pillars of our understanding. Two main lines of evidence link DA and D2Rs to SCZ. First, all effective antipsychotic treatments rely on pharmacological antagonism of D2Rs [6, 7]. Second, PET brain imaging studies consistently demonstrate increased dopamine release and elevated striatal D2R occupancy in SCZ [8] [9, 10]. Furthermore, (18)F-DOPA PET imaging found elevated striatal DA release capacity early on in adolescent subjects with high risk for schizophrenia [11] . While it is unknown how early changes in the dopamine system occur in patients with schizophrenia, Drd2 and other DA-related genes contain risk alleles that may affect brain development early on [12].

The postnatal development of the basal ganglia including the DA system is orchestrated by sensitive periods with enhanced neuroplasticity [13]. The closing of each sensitive period reduces this plasticity, after which the circuitry operates within narrower boundaries that shape its long-term function into adulthood. In rodents, striatal SPNs undergo a prolonged maturational process, in which hyperexcitable immature SPNs acquire their mature morphology, firing properties and synaptic input by the end of the fourth postnatal week [14, 15]. This process is triggered by the gradual rise in DA neurotransmission to SPNs and is dependent on Kir2 currents [16]. In parallel, early postnatal synaptic activity of maturing SPNs has been shown to regulate the formation of cortico-striatal synapses in a process that is likely to be influenced by DA signaling [17, 18].

Prior studies published by our group and others point to distinct effects of striatal D2R overexpression on incentive motivation depending on whether the manipulation occurs in adulthood or during development [19, 20]. Motivation is a fundamental brain function that allows individuals to select appropriate actions that lead to a goal by evaluating the benefits (value) of a reward and the cost (work) associated with obtaining it [21]. Impaired motivation is a symptom present in many neuropsychiatric disorders, including schizophrenia (SCZ) [22] and may arise from distorted computations of benefits and/or costs.

Virally induced D2R overexpression by indirect pathway neurons (iSPNs) in the NAc of adult mice increased motivation as reflected in enhanced performance in an operant Progressive Ratio (PR) task [23, 24]. This effect was shown to be mediated by decreased iSPN inhibitory output to the ventral pallidum (VP) [23]. However, the opposite effect on motivation was observed when D2R overexpression started from the late embryogenic period onwards (D2R-OE_dev_ mice) [25]. Here, a transgenic Tet-OFF system was used, resulting in diminished motivation in the PR task [25, 26]. In D2-OE_dev_ adult mice, NAc DA release and VTA neuronal activity are both reduced, suggesting that long-lasting circuit effects in the DA system may underlie these behavioral deficits [27–30]. These studies indicate that the developing indirect pathway plays an important role in the development of the adult DA system. However, in the experiments with D2-OE_dev_ mice overexpression of D2R occurred during the entire postnatal development. Here, we addressed the question whether inhibition of the indirect pathway during a defined postnatal time window can lead to long lasting changes in behavior and DA function.

To selectively target iSPNs during a defined postnatal time window, we expressed the designer receptor (DREADD) hM4Di in the indirect pathway of the striatum in neonates. Like D2Rs, hM4Di is a Gα_i_-and arrestin coupled GPCR that reduced neuronal excitability and inhibits synaptic transmission when activated by a synthetic ligand [31]. Using this approach, Kozorovitskiy et al developmentally inhibited the indirect pathway from postnatal day P8 to P15. The developmental inhibition increased the strengths of cortical synaptic input into the striatum and this increase could still be measured 10 days after the inhibition subsided [17]. However, whether early postnatal alterations in striatal circuit function leads to long-lasting changes in behavior that can be observed in adulthood remains to be shown. Here we show that developmental inhibition of iSPNs from P8-15 leads to long lasting decreases in open field activity and a deficit in progressive ratio performance. Moreover, using fiberphotometry in combination with a genetically encoded sensor for dopamine we found that NAc dopamine release does not adapt as readily to changes in effort requirements possibly explaining the decreased performance in the progressive ratio task. This establishes postnatal days P8-15 as a sensitive time window in the development of striatal circuitry as well as a period in which neural development can impact later adult motivated behavior.

## Methods

### Ethics compliance statement

All experimental procedures were conducted in accordance with ethical standards for scientific research stipulated by the NIH and Columbia University. All experimental procedures were approved by Institutional Animal Care and Use Committees by Columbia University and the New York State Psychiatric Institute (NYSPI protocol #1621).

### Animals

Homozygous Adora2a-Cre mice on a C57Bl6 background were crossed to C57Bl6 to obtain heterozygous Adora2a-Cre mice (RRID:MMRRC_036158-UCD), which were used in all experiments. Mice were housed 1–5 per cage for most experiments on a 12-hour light/dark cycle in a temperature-controlled environment (72^°^F and humidity 30-70%), with food and water ad libitum, unless otherwise noted. All experiments were conducted in the light cycle.

### Surgical procedures

#### Neonatal hM4D viral injection

On postnatal day 1 (P1) heterozygous Adora2a-Cre mouse neonates were injected bilaterally with the Cre-recombinase-dependent viral vectors AAV5-hSyn-DIO-hM4D-mCherry virus (Addgene Cat. No. 44362-AAV5-7x10^12^ vg/mL, Cambridge, MA) or AAV5-hSyn-DIO-GFP virus (Addgene Cat. No. 50457-AAV5, 7x10^12^vg/mL). The viral injection procedures were executed as described in [32]. Briefly, a custom-made neonatal stereotaxic adaptor was attached to a Digital stereotaxic apparatus for mice (Stoelting Company, Wood Dale, IL) equipped with a Nanoject II programmable nanoliter injector (Drummond Scientific Company, Broomall, PA). Each Adora2a-Cre P1 neonate was removed from the nest and placed in ice for 2 minutes to induce anesthesia. The anesthetized pups were then transferred to the stereotaxic frame and their heads secured to the head cradle with cloth tape in the neonatal device. The coordinates of the Nanoject II were then “zeroed” on the location of lambda, verifiable by visual inspection. For striatal targeting, the following stereotaxic coordinates were used (from vascular lambda): AP +2.50mm, ML ± 1.30mm, DV -3.45mm, -3.15mm and -2.30mm (measured from the surface of the skin). At each DV position, 50nl of virus were injected with 1 minute between each DV position. After removal of the injection needle, the procedure was repeated on the contralateral side. Anesthesia was maintained throughout the procedure by surrounding the pups with a small amount of ice chips. Each procedure was timed to last less than 20 minutes total. After completion of the procedure, pups were carefully removed from the stereotaxic frame and returned to the nest for recovery.

#### Adult dLight1.2 viral injections and optic fiber implantation

Adult mice (10-12 weeks) were anesthetized with 4% isoflurane and maintained at 1–2% throughout the entire procedure. Mice were unilaterally injected using a Nanoject II programmable nanoliter injector with 600 nL/hemisphere with AAV5-hSyn-dLight1.2 (Addgene Cat. #111068-AAV5) [33] into the NAc core using stereotaxic bregma-based coordinates: AP, +1.7 mm; ML, ±1.2 mm; DV, –4.1, –4.0, and –3.9 mm (200 nL/DV site). Each 200nL injection was subdivided into 40nL increments separated by 1-minute intervals. The on/off kinetics for the dLight1.2 sensor are 9.5 and 90 ms, respectively [34]. Following virus injection, 400 µm fiber optic cannulas (Doric, Quebec, Canada) were carefully lowered to a depth of –3.9 mm and fixed in place to the skull with dental cement anchored to machine mini-screws. Groups of mice used for experiments were housed in a counterbalanced fashion that accounted for sex, age, and home cage origin. Cannula-implanted mice began behavioral training 4 weeks after surgery. At the end of experiments, animals were perfused with 4% paraformaldehyde (PFA) and brains were processed post hoc to validate virus expression and optic fiber location as previously described [35, 36]

#### DREADD (hM4DGi) developmental activation with agonists

For the first mouse cohorts used in this study (Figs. 1 and 2), clozapine-n-oxide (CNO) dissolved in sterile 0.9% saline (1 mg/kg) was injected intraperitoneally during the postnatal period of P8-15 twice daily. Twice a day 0.9% saline i.p. injection were given to hM4D-expressing littermates as control, Mice expressing GFP as a control virus received the identical treatment as hM4D-expressing A2a-Cre mice. The dose and frequency of CNO injections were based on previous experiments and published data from our group [23, 37]. For a second cohort of mice used in the open field, PR experiments, and fiber photometry studies, the DREADD agonist JHU37160 (“J60”) was used. J60 was dissolved in 0.9% sterile saline and injected intraperitoneally twice daily at a dose of 0.1 mg/kg. This dose was derived from [38].

**Figure 1:**
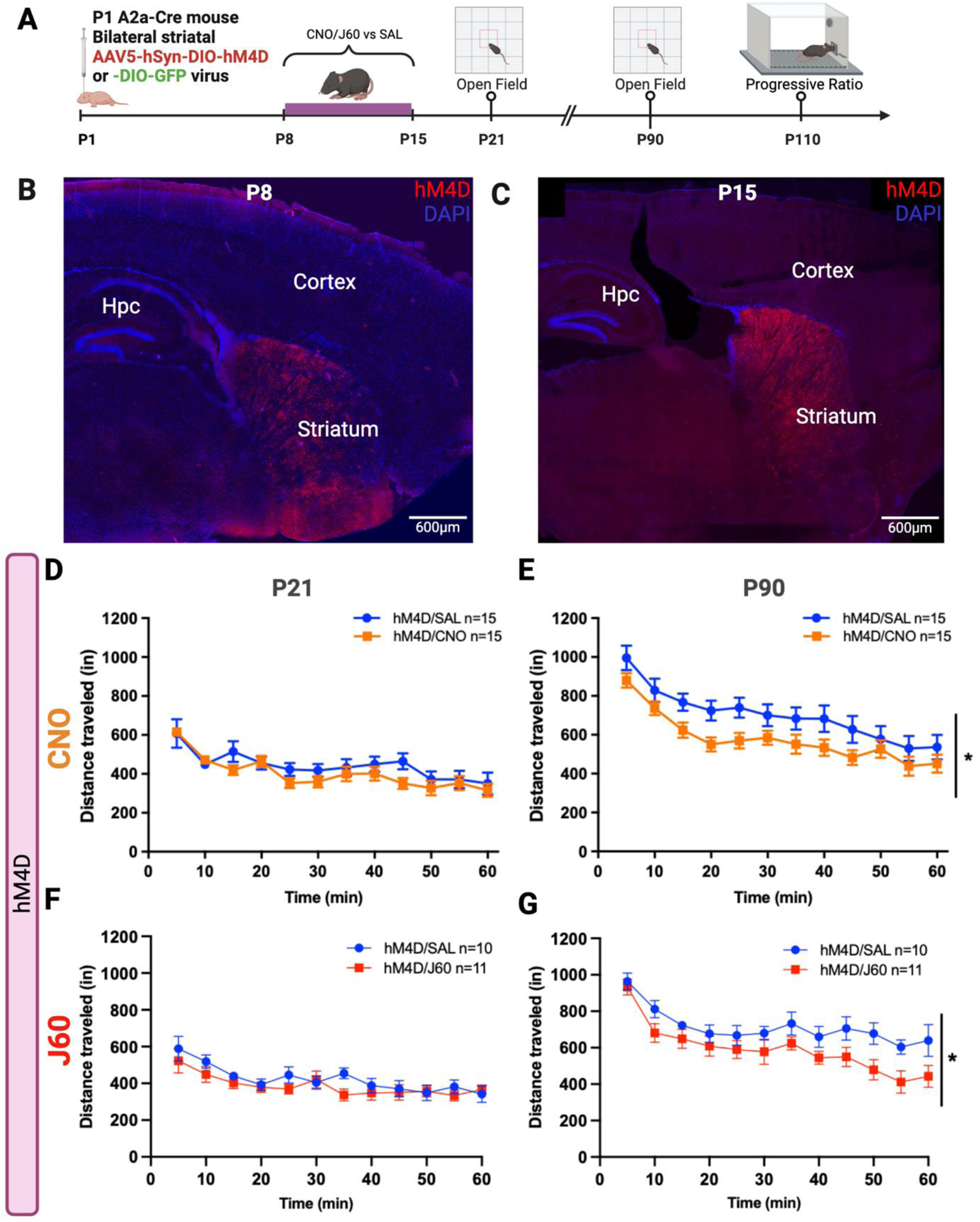
Selective neonatal inhibition of striatal indirect pathway leads to reduced Open Field locomotion. **A)** Experimental timeline from stereotaxic viral hM4D (or GFP) injection, DREADD agonist treatment (CNO or J60), to behavioral assessments. **B-C)** Representative confocal images of a sagittal section of a P8 **(B)** and P15 **(C)** A2a-Cre/hM4D^dev^ mouse showing expression of hM4D-mCherry in the striatum (Hpc: Hippocampus). **D-E)** Distance traveled by P21 **(D)** and P90 **(E)** old A2a/hM4D^dev^ mice after developmental injection of CNO or saline from P8- 15, respectively. *: RM 2-way ANOVA, Treatment Factor [F (1, 28) = 4.225, p=0.0493]. **F-G)** Distance travelled by P21 **(F)** and P90 **(G)** old A2a/hM4D^dev^ mice after developmental injection of J60 or saline from P8-15, respectively. *: RM 2-way ANOVA, Treatment Factor [F (1, 19) = 4.797, p=0.0412].

**Figure 2:**
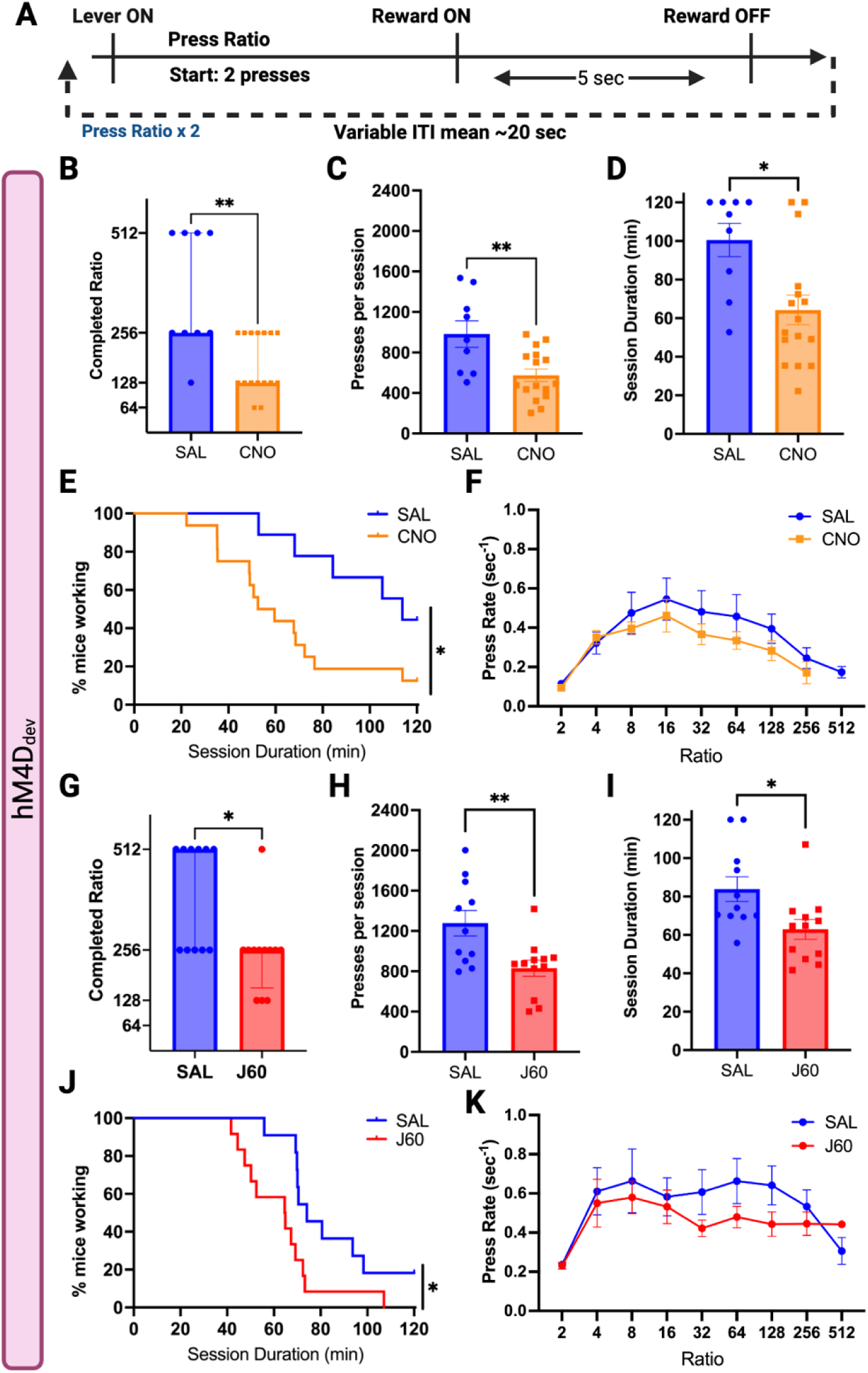
Developmentally inhibited A2a-Cre/hM4D^dev^ mice display reduced willingness to work for food. **A)** Schematic of Progressive Ratio (PR) trial structure (*ITI: inter-trial interval).* Sessions started with a ratio of 2 presses/reward and doubled after each completed trial. Sessions were timed out after 2 hours or if mice didn’t press for 5 minutes. **B-F)** PR results for CNO-treated A2a/hM4D^dev^ mice, n=9-16. **B)** Session breakpoints (earned rewards) for saline- and **CNO**-treated mice. Data expressed as median ± interquartile range, n=9-16, 2-tailed unpaired Mann-Whitney test, **p=0.0042. **C)** Summary of total presses per session for saline- and CNO-treated mice, expressed as mean ± SEM, 2-tailed unpaired t-test, **p=0.0039. **D)** Survival analysis of percentage (%) of mice engaged in task as a function of session duration. Mantel-Cox Log-rank test, *p=0.0234. **E)** Mean (± SEM) press rate (sec^-1^) as a function of ratio, Mixed-effects Analysis, Ratio effect (p<0.0001), Treatment effect (p = 0.2397). **F)** Mean (± SEM) latency to reward (sec) as function of ratio, Mixed-effects Analysis, Ratio effect (p=0.2750), Treatment effect (p=0.5973) **G-K)** PR results for **J60**-treated A2a/hM4D^dev^ mice. **G)** Effort breakpoints (completed ratio) for saline- and CNO-treated mice. Data expressed as median +/- interquartile range, n=11-12, 2- tailed unpaired Mann-Whitney test, *p=0.0104. **H)** Summary of total presses per session for saline- and CNO-treated mice, expressed as mean +/- SEM, n=11-12, 2-tailed unpaired t-test, ** p =0.0062. **I)** Summary of session duration, expressed as mean +/- SEM, n=11-12, 2-way unpaired t-test, *p=0.0185. **J)** Survival analysis of % of mice engaged in task as a function of session duration. Mantel-Cox Log-rank test, * p=0.0149. **F)** Mean (+/- SEM) press rate (sec^-1^) as a function of ratio, n=11-12, RM 2-way ANOVA.

### Behavioral training

#### Open Field

P21 and P90 A2a-Cre expressing hM4D (or GFP) and developmentally treated with CNO or J60 were tested in open field boxes (OF) equipped with infrared photobeams to measure locomotor activity (Med Associates, St. Albans, VT). Movement data were acquired using Kinder Scientific Motor Monitor software (Poway, CA) and expressed as total distance traveled over 60 minutes.

#### Progressive Ratio (PR) task

For all operant behavioral tasks (PR, FR, and SR), operant chambers (Med-Associates, St. Albans, VT) equipped with retractable levers, liquid dippers and kept in sound/light-attenuating cabinets were utilized. Infrared photocell sensors located in the reward magazine detected head entries during each behavioral trial. Each dipper delivered a drop (approximately 15μL) of evaporated milk. A “house light” illuminated each chamber during the tasks. Med-PC IV and V software were used to control and record data acquired by each operant chamber. Adult mice (> 90 days) were first trained to access the dipper, then to lever press and after went through progressive interval and ratio schedules. Mice were weighed daily and food restricted to 85-90% of baseline weight, while access to water remained unrestricted.

On day 1 of dipper training, mice were exposed to “free” (no lever press required) dipper presentations each lasting 8 seconds before being retracted. If mice entered the food magazine when the dipper was elevated (“dipper up”), the event was recorded as a rewarded head entry. Head entries that occurred when the dipper was not up were recorded as unrewarded head entries. Day 1 sessions were capped at a total of 20 trials or 30 minutes elapsed. Trials were separated by a variable inter-trial interval (ITI) of mean 20 seconds. On day 2 of dipper training, trials consisted of 5-second dipper presentations separated by variable ITI of mean 20 seconds. Sessions ended at after 45 trials (capped at 45 minutes total) and criterion was reached after 30 rewarded head entries.

For lever press training, a lever was extended at the beginning of a trial and a successful press was reinforced with a dipper presentation (for 5 seconds) on a continuous reinforcement schedule (CRF) with a variable ITI (mean 20 sec). Levers were retracted after each reward. Criterion was reached after 45 rewarded lever presses within a 60- minute session.

After CRF training, mice were trained on random interval (RI) training schedules, in which a lever press was rewarded after a variable time since the previous reward. Over successive days, the mean intervals were5, 10, 15, and 20 seconds (RI5, RI10, RI15, RI20) on consecutive days. RI20 was repeated for 3 consecutive days and on the following day, mice were exposed to the Progressive Ratio (PR) task, in which the ratio of presses to reinforcements doubled after each trial, starting with a ratio of 2 presses/reward, followed by 4, 8, 16, 32, 64, 128, 256, 512, 1024, 2048 until the session ended at 120 minutes or terminated after 5 minutes of inactivity (no lever presses). We based samples sizes on previous experiments and no statistical methods were used to calculate sample sizes [23].

### Fixed Ratio (FR) combined with fiber photometry

Four weeks after dLight1.2 viral delivery and optic fiber implantation, adult mice (>P90) were food restricted to 90% of their baseline weight and underwent dipper and CRF training as described above while tethered to an optical cable to be used for fiber photometry (FP) measurements. Mice were then trained on 3 levels of Fixed Ratio (FR): FR10, FR20, FR40, in which milk rewards were delivered after 10, 20, or 40 lever presses in a trial, with a fixed ITI of 10 seconds. Each FR requirement was repeated on three consecutive days (an FR “block”) and all sessions consisted of 45 trials (or 60 minutes maximum) Data from each FR block was pooled across days for each animal.

### Stepping Ratio (SR) combined with fiber photometry

Adult mice (>P90) expressing dLight1.2 and implanted with an optic fiber in the NAc were food restricted to 90% of their baseline weight and underwent dipper and CRF training while tethered to a fiber optic cable, as described above. After CRF training, mice were trained on RI schedules starting with RI5, followed by RI10, RI15, and RI20 (x 3 days). Next mice were exposed to a stepwise progressive ratio schedule, in which each ratio was repeated for 10 trials before doubling to the next ratio. The ratios were 25, 50, 100, 200, and 400. The ITI was variable with mean of 20 seconds. Sessions were kept at a 90-minute maximum.

### In vivo fiber photometry

Fiber photometry equipment was set up and operated as described in [36]. Briefly, the equipment consisted of a 4-channel LED driver (Doric) connected to 2 sets of a 405-nm and a 465-nm LEDs (Doric). The 405-nm LEDs were filtered through 405-410 nm bandpass filters, whereas 465-nm LEDs passed through 460-490 nm GFP excitation filter using two 6-port Doric minicubes. Dichroic mirrors were connected to each 405 and 465 nm pairs to split emission and excitation lights. Fluorescent signal from the mouse’s implanted optic fiber cannula was transmitted by low autofluorescence optical patch cords (400μm/0.48 NA, Doric) connected to the minicubes and filtered through 500-540 nm GFP emission filters coupled to photodetectors with the gain set to “DC Low” (Doric). Signals were sinusoidally modulated with Synapse software and RZ5P Multi I/O Processors (Tucker-Davis Technologies), at 210 Hz (405 nm) and 330 Hz (465 nm) and low-pass filtered at 3Hz via a lock-in amplification detector. Power at the patch cord for 405 and 465nm was set at 30μW or below.

Custom in-house MATLAB scripts were used to process fiber photometry (FP) and behavioral data. Operant behavioral output was time-aligned to the FP signal by a cable that, each time the dipper mechanism was activated, sent a square pulse from the operant chamber to the FP acquisition system. The alignment of the FP signal with other behaviorally salient events (i.e. lever extension, lever presses, head entries) was derived from the alignment with the dipper activation events. The signal from the 405nm channel was used to control for movement/noise artifacts and 465nm channel was used to detect the main fluorescent signal emitted by dLight1.2 upon DA binding. The demodulated signals were extracted in 15 sec windows surrounding each event, which was determined as time = 0. Signals (405 and 465nm) were down sampled by a factor of 10 using a moving window mean. The change in fluorescence, ΔF/F (%), was defined as (F-F_0_)/F_0_ x 10, where F is the fluorescent signal (465nm channel) at each time point, whereas F_0_ was calculated by applying a least-squares linear fit to the 405nm signal to align with the 465 nm signal [39]. Normalization of signals across animals and sessions for each behavioral event was achieved by calculating a local fluorescence baseline value for each trial using the average of the preceding 5 sec and subtracting that value from the signal. The daily average dLight1.2 signal traces were calculated using session average traces from individual mice. Dopamine peak amplitudes were calculated as the maximal change of the signal from the local baseline. The area under the curve (AUC) was calculated as the total net area between Y = 0 and the session averaged signal trace for defined time intervals for each epoch evaluated: Lever extension (0 sec – 1.5 sec) and Dipper up (0 sec – 1.2 sec).

### Histology

Adult mice were anesthetized with an i.p. injection of 100mg/kg ketamine and 5mg/kg xylazine prior to being perfused with 4% paraformaldehyde (4% PFA) in PBS. Brains were post-fixed in 4% PFA overnight before being transferred to 0.5% PFA for long-term storage. Brains were sectioned at 40μm on a vibratome (Leica, Buffalo Grove, IL, USA). The following primary antibodies were used: mCherry (rabbit-anti-DsRed; Takara Bio, Mountainview, CA, USA; 632496, 1:250) or green fluorescent protein (chicken anti-GFP; Abcam, Cambridge, UK, ab13970, 1:1000). Primary antibody incubation was 16h at 4°C. Alexa Fluor-conjugated secondary antibodies (donkey anti-rabbit Alexa Fluor-546 and goat anti-chicken Alexa Fluor-488, Invitrogen, 1:1000) were used for fluorescent signal detection and incubated for 1 hour at room temperature. Stained tissue slices were then mounted on glass slides with Vectashield containing DAPI (Vector Labs). Viral expression was confirmed from mCherry or GFP staining, and locations of recording site lesions were confirmed under DAPI under an epifluorescence microscope (Zeiss).

For P8-15 immunostaining, fixed pup brains were imbedded in an Optimal Cutting Temperature (OTC) medium, stored under -20°C, and sectioned on a cryostat (Leica, Buffalo Grove, IL, USA) at a thickness of 20 μm. Brain sections were transferred to positively charged glass histology slides (Fisherbrand) and immunostained with mCherry (rabbit-anti-DsRed; Takara Bio, Mountainview, CA, USA; 632496, 1:250) directly on the histology slide. Sections were then mounted with DAPI-containing Vectashield mounting medium and imaged on a confocal microscope (Leica).

### Automated cell counting

Automated segmentation and counting of immunostained Cre-positive cells was achieved in anatomically defined areas (410.67μm x 410.67μm) using an ImageJ/FIJI workflow based on Trainable Weka Segmentation [40]. Cell positive for mCherry were manually scored by two human subjects blind to the experimental conditions. Colocalization was defined by centroid-based proximity between independently detected Cre and mCherry signals across six sections (N=2 mice).

### Statistical Analysis

Operant behavioral data were analyzed using custom made MATLAB (MathWorks) scripts. Statistical analyses were performed on GraphPad Prism 9 (GraphPad) and MATLAB (MathWorks). Data were expressed as mean ± standard error of the mean (SEM), except where noted. Unpaired 2-tailed Student’s t-test was used compare the mean between two groups for normally distributed data. For comparing two non-Gaussian data sets the Mann-Whitney test was used. Full Model (including interaction term) 2-way Repeated Measures (RM) ANOVA, with Geiser-Greenhouse correction for sphericity, was used to compare two or more groups over a period of time or press ratio. Sidak correction was used for multiple comparisons. In the rare instances of missing values in repeated measures analysis, a mixed effects model was used as implemented in Prism 9. For comparing survival curves the Log-Rank (Mantel-Cox) test was used. For correlational analysis of photometry and behavioral data, Spearman correlation coefficients were used. Statistical significance was set at p-value < 0.05. Both males and females were analyzed, but the sample sizes were not powered to detect sex differences. Investigators were blinded to treatment (CNO, J60, or SAL) throughout experimentation and analysis.

## Results

### Inhibition of indirect pathway during the second postnatal week decreases locomotor activity in adult A2a-Cre/hM4D_dev_ mice

In a first experiment, we determined whether inhibition of the indirect pathway from P8-15 affects locomotor activity in the open field hile mice were juveniles and then again as adults (for timeline of experiment see **Fig. 1A**). To inhibit the indirect pathway from P8-15 we injected a Cre-dependent adeno-associated virus (AAV) carrying the inhibitory DREADD, hM4DGi, into the striatum of P1 Adora2a-Cre neonates, or a GFP virus as control. The bilateral neonatal viral injections were accomplished by using a mouse neonatal stereotaxic adaptor developed by our group, which enhances reliability of injections [32]. P1 injection leads to robust expression at P7 (**Fig. 1B**) with a further increase in expression by P15 (**Fig. 1C**). We estimated the transfection efficacy in P15 mice and found that approximately 44.49 ± 6.46 % (mean ± SEM) of Cre recombinase-positive (Cre+) striatal cells co-stained for hM4D-mCherry throughout the striatum (**Supplementary Fig. S1**). Half of the mice were injected i.p. with the DREADD agonist clozapine-n-oxide (CNO; 1 mg/kg) and half with saline, twice per day.

We first tested the mice in the open field since developmental D2R-OE mice showed a decrease in open field performance [26]. A2a/hM4D_dev_ mice were introduced to the open field arena during infancy (P21) and adulthood (P90) for 60 minutes. When tested on P21, P8-15 CNO-treated mice did not display differences in locomotion (distance traveled) when compared to control mice (Repeated Measures 2-way ANOVA, n=15, Drug factor [F (1, 28) = 1.187, p=0.2852], Time factor [F (6.575, 184.1) = 12.47, p<0.0001], Drug x Time interaction [F (11, 308) = 0.9838, p=0.4610], **Fig. 1D**). On P90, the same mice were placed in the open field arena and an overall decrease in locomotion emerged (RM 2-way ANOVA, n=15, Treatment factor [F (1, 28) = 4.225, p=0.0493], Time factor [F (5.377, 150.6) = 32.52, p<0.0001], Treatment x Time interaction [F (11, 308) = 0.7023, p=0.7363], **Fig. 1E**).

To control for potential non-specific effects of CNO due to its conversion to clozapine [41] a separate cohort of animals received a Cre-dependent GFP virus at P1 and treated with either CNO or saline during the same P8-15 period. RM 2-way ANOVA analyses found no differences in OF distance traveled at either P21 (Treatment factor [F (1, 16) = 0.7182, p=0.4092], Time factor [F (5.411, 86.57) = 7.337, p<0.0001], Treatment x Time Interaction [F (11, 176) = 1.173, p=0.3089]) or P90 (Treatment Factor [F (1, 15) = 0.8230, p=0.3787], Time Factor [F (5.528, 80.40) = 23.75, p<0.0001], Treatment x Time Interaction [F (11, 160) = 0.6359, p=0.7961, Supplementary Fig. S2).

We then attempted to replicate the results using the recently developed hM4D agonist, JHU37160 (J60), that is not metabolized to clozapine [38]. To activate hM4D_Gi_ J60 was injected systemically twice daily at 0.1mg/kg from P8-15. Similar to the CNO-treated mice we observed decreased locomotor activity emerging at P90 (RM 2-way ANOVA, n=10-11, Time factor [F (4.635, 88.07) = 17.44, P<0.0001], Treatment factor [F (1, 19) = 4.797, p=0.0412], Treatment x Time interaction [F (11, 209) = 1.036, p=0.4157]; Fig. 1G) but not at P21 (RM 2-way ANOVA, n=10-11, Time factor [F (3.815, 72.48) = 6.406, p=0.0002], Treatment factor [F (1, 19) = 1.295, p=0.2692], Treatment x Time interaction [F (11, 209) = 0.7808, p=0.6591], **Fig. 1F**).

A separate cohort had a Cre-dependent GFP virus expressed neonatally and treated with either J60 or saline from P8 to P15, followed by OF testing at P90 showed no differences in distance traveled (RM 2-way ANOVA, n=9, Time Factor [F(5.173, 82.76)=27.37, p<0.0001], Treatment (J60) Factor [F(1,16)=0.3312, p=0.5730], Time x Treatment interaction [F(11,176)=0.9033, p=0.5384, **Supp. Fig. S2**).

Together, these results show that the P8-15 inhibition of indirect pathway leads to a decrease in locomotor activity that manifests in the adult animal. As we tested male and female mice in both the CNO and the J60 experiments we tested for sex effects. However, we no sex x treatment interaction was measured on open field performance (data not shown).

### Inhibition of the indirect pathway during the second postnatal week decreases motivation in adult A2a-Cre/hM4D_dev_ mice

To assess whether the developmental inhibition leads to long lasting changes in reward driven motivated behavior, we tested A2a/hM4D_dev_ mice in the Progressive Ratio (PR) operant task at P110. Briefly, adult mice were trained on a continuous reinforcement schedule (CRF) and progressively longer random interval schedules as described in the method section. This was followed by a PR day, during which the press-to-reward ratio doubled with each trial, starting with 2 presses/reward, quickly reaching 2048 presses/reward after 11 trials. Sessions were automatically timed out at 2 hours or if the mice did not press for 5 minutes (**Fig. 2A**). The final ratio reached by each animal was defined as the breakpoint.

We evaluated a cohort of A2a/hM4D_dev_ mice treated with CNO (1mg/kg) from P8-15 (developmental inhibition), twice daily versus saline-treated A2a/hM4D_dev_ mice at age P90-110 (**Fig. 2B-F**). We analyzed the following PR outcomes: breakpoint, total number of lever presses, session duration (up to 120 minutes), session “survival” analysis, press rate for each of the ratio requirements, and the latency to reach the reward magazine (latency to reward) for each ratio. Compared to the control group (median total rewards = 8 [completed ratio of 256 presses/reward], n=9, **Fig. 2B**), developmentally inhibited (CNO) mice displayed a reduced mean ratio breakpoint (median total rewards = 7 [completed ratio of 128 presses/reward], n=16, 2-tailed unpaired Mann-Whitney test, U=24.5, p=0.0042, **Fig. 2B**). The non-parametric Mann-Whitney test was chosen because the breakpoint data were not normally distributed. CNO-treated mice also showed significantly lower mean total lever presses (575.1 ± 61.85 presses), relative to saline controls (982.8 ± 130.1 presses, 2-tailed unpaired t-test, n=9-16, t=3.210, df=23, p=0.0039, **Fig. 2C**). The analysis of the mean session duration revealed that on average CNO-treated mice had more abbreviated sessions (34.2% shorter) than control mice (CNO: 66.17 ± 7.91 minutes, n=9; SAL: 100.5 ± 8.6 minutes, n=16; 2-way unpaired t-test, t=2.811, df=22, p=0.0102). The session “survival” analysis confirmed that the CNO-treated mice (median survival = 56.04 minutes) quit the PR task at earlier time points, while the control group (median survival = 113.8 minutes) maintained task engagement for longer periods (Mantel-Cox Log-rank test, χ^2^ = 5.139, df=1, p=0.0234, **Fig. 2D**).

The press rates by both groups were sensitive to the progressively higher ratios in inverted “U-shaped” response curves, peaking at the ratio of 16 (**Fig. 2E**). Although the CNO group’s response curve appears slightly lower than the control group, there was no statistically significant effect of treatment (Mixed-effects Analysis, n=9-16, Ratio Fixed Effect [F(2.552, 47.53) = 22.75, p <0.0001], Treatment Fixed Effect [F(1, 20) = 1.469, p=0.2397], **Fig. 2E**). Lastly, the Mixed-effects analysis of the latency to reward revealed no effect of treatment or ratio requirements (Mixed-effects Analysis, n=9-16, Ratio Effect [F(2.392, 47.24) = 1.333), p = 0.2750], Treatment Fixed Effect [F(1, 23) = 0.2870, p = 0.5973], **Fig. 2F**).

To test for possible effects of hM4D-mediated effects of CNO alone, an independent cohort of A2a-Cre mice were injected with an AAV5-hSyn-DIO-GFP virus bilaterally in the striatum at P1 (A2a/GFP), then followed by twice daily i.p. CNO injections (1mg/kg) or normal sterile saline from P8-15 (**Supp. Fig. S3**). At P90-110, the mice were tested on the same PR protocol as described above. No significant differences were detected between A2a/GFP^CNO^ and A2a/GFP^SAL^ mice on breakpoint (GFP^CNO^: median rewards = 8 [completed ratio = 256], GFP^SAL^: median rewards = 8, n=10-11, 2-way unpaired Mann-Whitney test, U=48.5, p=0.5763, **Supp**. **Fig. S3A**), mean total presses (GFP^CNO^: 728.8 ± 77.44, GFP^SAL^: 743.1 ± 78.98, n=10-11, 2-way unpaired t-test, t=0.1288, df=19, p=0.8989, **Supp. Fig. S3B**), mean session duration (GFP^CNO^: 89.89 11.17 minutes, GFP^SAL^: 102.1 ± 6.55 minutes, n=10-11, 2-way unpaired t-test, t=0.9636, df=19, p=0.3473, session “survival” (GFP^CNO^: median 95.79 minutes, GFP^SAL^: 105.6 minutes, Mantel-Cox Log-rank test, χ^2^ = 0.1838, df=1, p=0.6681, **Supp. Fig. S3C**), press rate as a function of ratio (Mixed-effects, n=10-11, Ratio effect [F(2.422, 41.17) = 15.43, p<0.0001], Treatment effect [F(1,19) = 0.01908, p=0.8916], **Supp. Fig. S3D**), or latency to reach reward (Mixed-effects Analysis, n = 10-11, Ratio Effect [F (4.095, 67.57) = 5.864, p = 0.0004], Treatment Effect [F (1, 19) = 0.02145, p = 0.8851], **Supp. Fig S3E**).

Next, we evaluated a separate, independent cohort of A2a/hM4D_dev_ mice treated with J60 (P8-15, twice daily, 0.1mg/kg) or saline. Compared to saline controls (median rewards = 9, corresponding to a completed ratio of 512), the J60-treated group (median rewards = 8, corresponding to a completed ratio of 256) showed a significantly reduced breakpoint, (2-tailed unpaired Mann-Whitney test, n=11-12, U=28, p=0.0104, **Fig. 2G**). The same was observed in the total number of presses, where J60-treated displayed reduced overall pressing compared to controls (830.4 ± 80.6 and 1278 ± 126.2 presses, respectively; 2-tailed unpaired t-test, n=11-12, t=3.038, df=21, p =0.0062, **Fig. 2H**). Moreover, an analysis of the mean session durations showed that J60-treated mice stopped pressing earlier than controls (62.9 ± 5.2 vs 83.9 ± 6.5 minutes in session, respectively; 2-tailed unpaired t-test, t=2.554, df=21, p=0.0185). A complementary “survival” analysis confirmed that J60-treated mice disengaged (defined as no presses for 5 minutes) from the task earlier than controls (Mantel Cox Log-rank test, χ^2^= 5.934, df=1, p=0.0149; **Fig. 2I**).

For both control and J60-treated A2a/hM4D^dev^ mice press rates varied in response to increasing ratios, but no statistical differences were detected between treatment groups (Mixed-effects Analysis, n=10-11, Ratio Fixed Effect [F (3.355, 59.13) = 7.003, p=0.0003], Treatment Fixed Effect [F (1, 20) = 0.6637, p=0.4248], **Fig. 2J**). The latency to reward analysis revealed no statistical treatment group or ratio differences(Mixed-effects Analysis, 10-11, Ratio Fixed Effect [F (3.297, 55.63) = 1.076, p = 0.3705], Treatment Fixed Effect [F (1, 20) = 0.3699, p = 0.5499], Ratio x Treatment [F (8, 135) = 1.243, P = 0.2792], **Fig. 2K**).

We also ran a control experiment to test for J60 long-term effects on PR performance. Like for CNO, A2a/GFP^dev^ control mice treated with J60 or saline during development showed no treatment differences with respect to session breakpoint (GFP^SAL^: median rewards = 9 [completed ratio of 512], GFP^J60^: median rewards = 9 [completed ratio of 512], n=7, 2-tailed unpaired Mann-Whitney test, U=19.5, p=0.6999, **Supp. Fig. S3F**), total press number (GFP^SAL^: 1986.0 ± 362.4 presses, GFP^J60^: 1657 ± 269.2 presses, n=7, 2-tailed unpaired t-test, t=0.7294, df=12, p=0.4797, **Supp. Fig. S3G**), mean session duration (GFP^SAL^: 83.87 ± 15.0 minutes, GFP^J60^: 76.26 ± 16.21 minutes, n=7, 2-tailed unpaired t-test, t=0.3446, df=12, p=0.7364), session survival (Mantel-Cox Log-rank test, χ^2^ = 0.0340, df=1, p=0.8537, **Supp. Fig. S3H**), press rate as function of ratio (Mixed-effects, n=7, Ratio effect [F(3.315, 35.80) = 5.559, p=0.0024], Treatment effect [F(1, 12) = 0.1663, p=0.6906], Interaction [F(10, 108)=1.824, p=0.0646], **Supp. Fig. S3I**), or latency to reward as function of ratio (Mixed-effects, n=7, Ratio effect [F(2.363, 24.16) = 1.862, p=0.1719], Treatment effect [F(1, 12) = 1.551, p=0.2368], Interaction [F(9, 92) = 1.294, p=0.2511], **Supp. Fig. S3J)**.

Together, these results show that the P8-15 inhibition of indirect pathway leads to an impairment in PR performance in adulthood. As in the open field experiments, we tested male and female mice but detected no sex x treatment interaction (data not shown).

### Developmental indirect pathway inhibition leads to reduced NAc dopamine release when effort requirements increase over days

Dopamine release in the NAc and VTA neuronal activity have both been shown to be reduced in the D2R overexpressing mice, that overexpresses D2Rs from early development on. This decrease in VTA/NAc dopamine function was associated with a decrease in PR performance suggesting that a long-lasting reduction in dopamine function may underlie the motivational deficits observed in D2R overexpressing mice [28, 42]. Like D2R overexpressing mice, A2a/hM4Ddev mice display reduced PR performance; therefore, we hypothesized that diminished DA release associated with increasing effort requirements may impose a limit on the effort the mice are willing to spent (i.e. lever pressing, reward seeking behavior, time to complete trials) in A2a/hM4D_dev_ mice.

The genetically encoded dopamine sensor, dLight1.2, was virally expressed in the NAc of A2a/hM4Ddev mice at P90, along with an optic fiber implant. Four weeks later mice were trained in the operant boxes while tethered to a fiber photometry (FP) acquisition system (**Fig. 3A**). Upon binding of DA, dLight1.2 receptors undergo a conformational change that activates a circularly permutated GFP protein, which emits a fluorescent signal reflecting extracellular DA levels [34]. Task-evoked DA transients were obtained by aligning the fluorescent signal to lever extension or reward presentation (see Methods).

**Figure 3.**
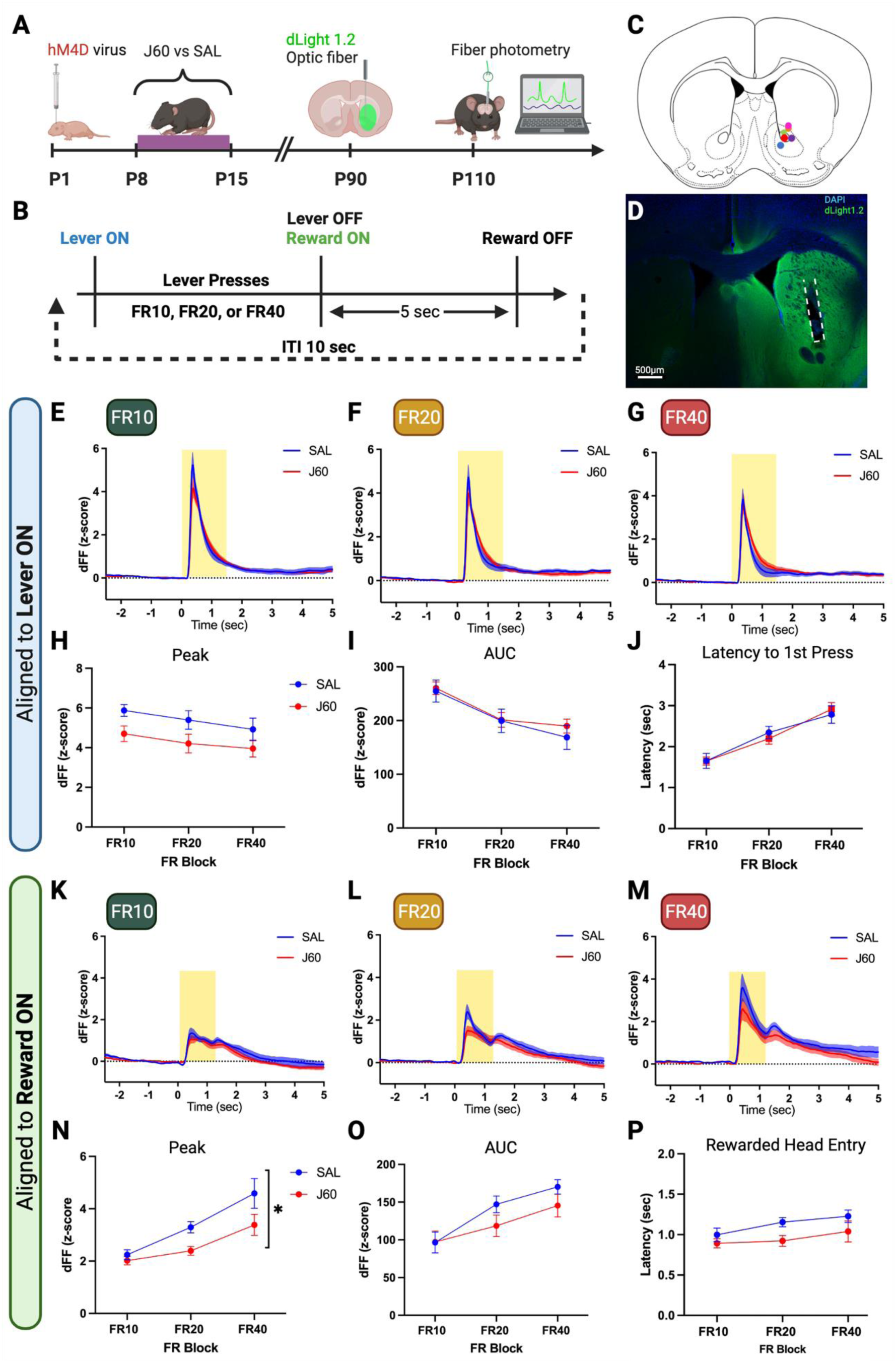
Developmentally inhibited mice show effort-sensitive reduction of task-evoked NAc DA release across several days. **A)** Experimental timeline from neonatal hM4D viral injection, developmental inhibition with J60 injections (P8-15), dLight1.2 viral injections and optic fiber implantation (P90), to simultaneous DA monitoring (fiber photometry) during operant tasks. **B)** Schematic of operant Fixed Ratio (FR) task structure. Each FR block was repeated for 3 consecutive days (45 trials per session, 135 total trials). A fixed ITI (inter-trial interval) of 10 seconds was used. **C)** Anatomical localization of optic fibers. **D)** Representative coronal section of viral dLight1.2 expression in striatum (anti-GFP immunohistochemistry) and optic fiber location (dashed lines delineate fiber shaft). **E-J)** Results of DA release aligned to lever extension (Lever ON; time = 0). **E-F)** Mean (+/-SEM) DA release (z-scored dFF) aligned to lever extension during FR10, FR20, and FR40, respectively (n=10). Yellow box represents time period used to calculate the area-under-the-curve (AUC). **H)** Mean peak amplitude (+/-SEM) of DA release (z-score dFF) across FR10, FR20, and FR40 shows declining peaks as work requirement increases for both groups (RM 2-way ANOVA, n =10, Ratio factor [F(1.682, 30.27) = 10.69, p=0.0006]), but only a statistical trend for treatment effect (Treatment factor [F (1, 18)= 3.43, p=0.0805]). **I)** Mean AUC (+/-SEM) from time 0 (lever ON) to 1.5s under FR10, FR20, and FR40 conditions. Both groups are sensitive to increasing work requirements (Ratio factor [F(1.736, 34.73)=30.75, p<0.0001]), but no effect of treatment or interaction. **J)** Mean (+/-SEM) latency to first press across FR10, FR20, and FR40 conditions showing both groups increasing latency with greater effort (Ratio factor [F (1.654, 31.42) = 33.84, p<0.0001]), but no effect of treatment or interaction. **K-P)** Summary results of DA release aligned to reinforcement delivery (Reward ON). **K-M)** Mean (+/-SEM) DA release (z-scored dFF) aligned to Reward ON during FR10, FR20, and FR40, respectively (n=10). **N)** Mean peak amplitude (+/-SEM) of DA release (z-score dFF) across FR10, FR20, and FR40 shows increasing peaks as work requirement increases for both groups (RM 2-way ANOVA, n=10, Ratio factor [F(1.159, 20.86) = 20.53, p=0.0001]), but a blunted rise in peak amplitude for the J60 group (Treatment factor F(1, 18) = 6.869, *p=0.0173). **O)** Mean (+/-SEM) AUC from time 0 to 1.2s under the three FR conditions shows the same positive relationship with increasing effort for both groups but no effect of treatment or interaction (RM 2-way ANOVA, n=10, Ratio [F(1.682, 30.27) = 24.83, p=0.0001]). **P)** Mean (+/-SEM) latency for rewarded head entries across FR blocks showing a mild positive relationship with effort, but no effect of treatment or interaction (FR block: F(1.501, 28.53) = 4.321, p=0.0323).

We first trained mice in a series of Fixed Ratio (FR) experiments starting from a ratio of 10 presses/reward (FR10), followed by FR20 and FR40. Each FR block repeated for 3 consecutive days, 45 trials each (**Fig. 3B**). At each trial initiation the lever is extended, which elicits a sharp DA (dLight1.2) fluorescent transient in both J60 and saline groups since lever extension predicts food reward (**Fig. 3E-G**). With each increase in the fixed ratio, the amplitude (peak) of these transients became reduced (RM 2-way ANOVA, n =10, Ratio factor [F(1.682, 30.27) = 10.69, p=0.0006], Fig. 3H). Our analysis also detected a statistical trend for effect of treatment (n =10, F (1, 18)= 3.43, p=0.0805, Fig. 3H), but no interaction effect between early inhibition and FR block (F (2, 36) = 0.2267, p=0.7983, Fig. 3H). The analysis of AUC DA transients showed a similar relationship between lever extension DA release and ratio (RM 2-way ANOVA, n =10, Ratio factor F(1.736, 34.73)=30.75, p<0.0001), but no effect of treatment (F(1, 20)= 0.1953, p=0.6632) or interaction between factors (F(2, 40)=0.4821, p=0.6210, Fig 3I). We examined the latency to start pressing the lever (latency to 1^st^ press) as a function of increasing effort. Both groups displayed longer latencies with higher effort ratios, but there was no effect of treatment or effort-treatment interaction (RM 2-way ANOVA, n=10, Ratio factor [F (1.654, 31.42) = 33.84, p<0.0001], Treatment factor [F (1, 19) = 0.004594, p=0.9467], Interaction [F (2, 38) = 0.4556, p=0.6375], Fig. 3J).

Next, we analyzed DA release aligned to reward availability cue (“dipper up”). In each trial, the dipper is only extended after the mouse completes the required number of presses for each ratio (10, 20, or 40). The DA transient aligned to “dipper” shows a biphasic structure, the first in response to dipper presentation, signaling reward availability, and the second, that is most likely related to the reward consumption (**Fig. 3K-M**). Opposite to lever extension, the DA transient aligned to the “dipper up” increased as the FR progresses from 10 to 40 presses/reward, while the second peak corresponding to reward consumption remains relatively stable across the changing FRs (**Fig. 3K-M**). Although both J60 (developmentally inhibited) and control groups exhibited an upward trajectory for the first DA transient, this response by J60-treated mice appeared to be blunted compared to control mice (**Fig. 3N-O**). A RM 2-way ANOVA of the “first peak” amplitudes across ratios was statistically significant for FR block and treatment, but not interaction (“FR block”: F(1.159, 20.86) = 20.53, p=0.0001; “Treatment”: F(1, 18) = 6.869, p=0.0173; “Interaction”: F(2, 36) = 1.477, p=0.2417; n=10, Fig. 3N). A RM 2-way ANOVA of the AUC corresponding to the first DA transient (Fig. 3K, yellow box) across all FR blocks was significant only for FR block (“FR block”: F(1.682, 30.27) = 24.83, p=0.0001; “Treatment”: F(1, 18) = 1.140, p=0.2997; “Interaction”: F (2, 36) = 1.696, p=0.1978; n=10, Fig. 3O). Thus, while in control mice DA release to a cue signaling reward availability is calibrated in response to increasing effort invested in the task, this calibration seems to be blunted in developmentally inhibited mice.

The increase DA peak responses to reward availability (the “first peak”) with increasing requirements, could be behaviorally reflected in the latency to reach the reward magazine with higher DA release associated with a faster head entry. However, head entry latencies increased with each increasing ratio. There was a statistical trend for the developmentally inhibited group (J60) reaching the magazine faster than controls, as well as an overall effect of effort (FR block) on reward latency, but no interaction effect (RM 2-way ANOVA, n=10, “Treatment” F(1, 19) = 3.109, p=0.0939; “FR block” F(1.501, 28.53) = 4.321, p=0.0323; “Interaction” F(F (2, 38) = 0.5149, p=0.6017; Fig. 3P).

### Developmental indirect pathway inhibition leads to reduced NAc dopamine release when effort requirements increase within the same day (Stepping Ratio)

During the PR task, mice experience an exponential increase in the effort requirement in each successive trial within the same session, forcing them to quickly adapt their behavior to the changing contingency. In FR series described above, mice experience increases in effort requirement at a much slower rate, with up to 135 repeated trials per fixed ratio, and the increase in ratio changing over days. Therefore, in each task information learned from a previous trial may have distinct behavioral effects on a subsequent trial which might also be reflected on dopamine release throughout the task [43].

Since the measurement of DA transients benefits from multiple trials for signal averaging, we designed a compromise with not too many trials that allow for slow adaptation but enough trials to average the fluorescent signal, while still permitting a sharp increase in effort requirement within the same session. In this Stepping Ratio task (SR), mice repeat each ratio for 10 trials before moving to the next one in the same session. Starting at a ratio of 25 presses/reward, each new ratio doubles after 10 trials (**Fig. 4A-B**).

**Figure 4.**
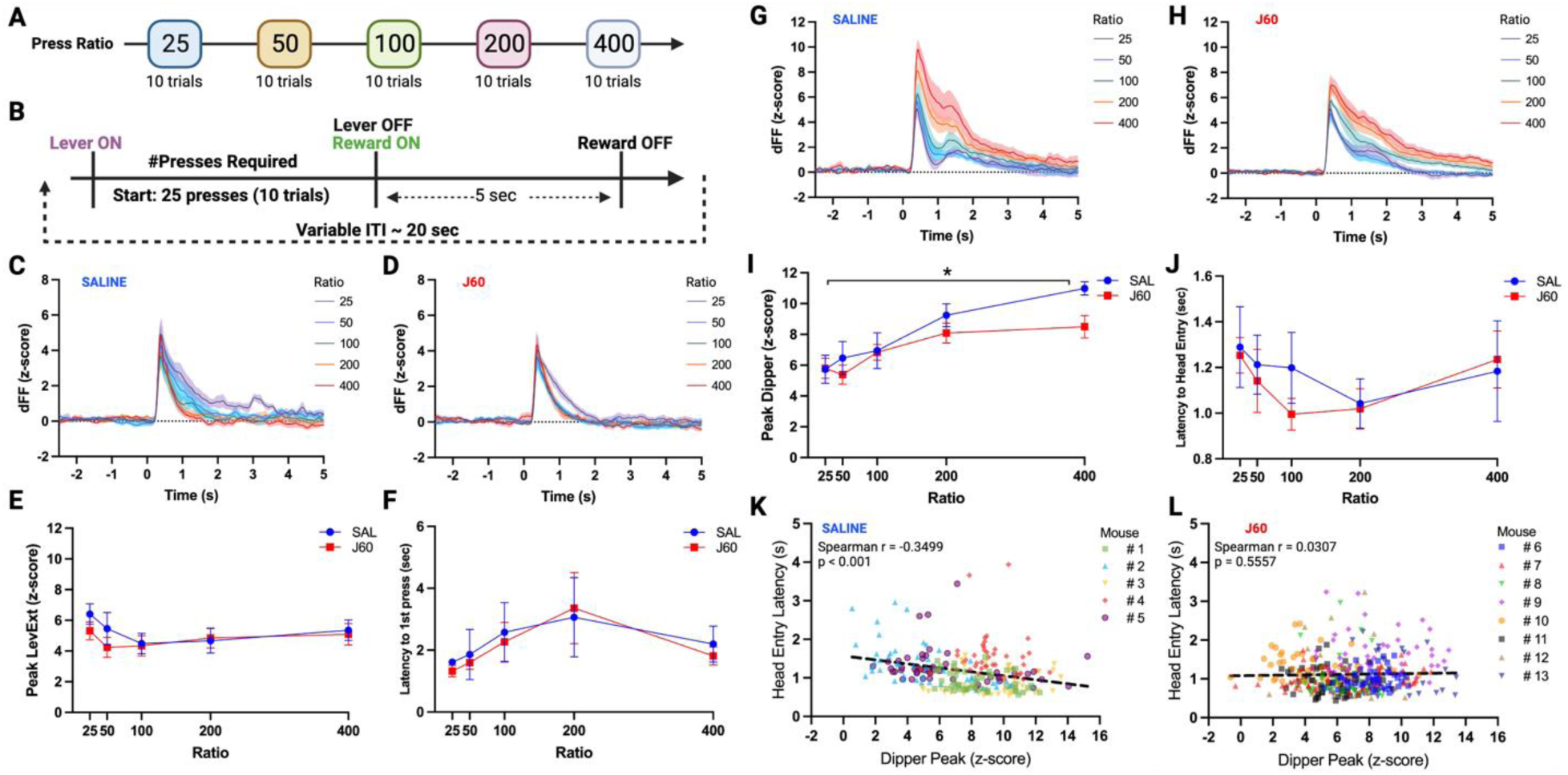
Developmentally inhibited mice have less adaptable reward-aligned DA release in response to rapidly increasing effort requirement. **A)** PR^dLight^ session structure. Each ratio was repeated for 10 trials before advancing to the next ratio. **B)** Schematic of PR^dLight^ trial structure. **C-D)** Mean (+/-SEM) DA release (z-scored dFF) aligned to lever ON (time 0s) across ratios 25 to 400 for saline and J60 groups, respectively. **E)** Mean (+/-SEM) peak DA release as a function of effort requirement, showing an overall effect of effort but not of treatment or interaction (RM 2-way ANOVA, n = 5-8, Ratio Factor [F (2.614, 28.76) = 3.742, p = 0.0263]). **F)** Mean (+/-SEM) latency to first press as function of increasing effort showing an inverted U overall relationship curve, but no statistical significance. **G-H)** Mean (+/-SEM) DA release (z-scored dFF) aligned to reward delivery (Reward ON = time 0s) across ratios 25 through 400 for saline (n=5) and J60 (n=8) groups, respectively. **I)** Mean (+/-SEM) peak DA release as function of ratio, showing blunted DA responses as effort increases (RM 2-way ANOVA, n= 5-8, Treatment x Ratio[F(4, 44) = 3.375, p = 0.0171]). **J)** Mean (+/-SEM) latency to head entry at each ratio; RM 2-way ANOVA did not show statistical significance for ratio, treatment or interaction. **K-L)** Scatter plots and Spearman correlations analyses for head entry latency and reward-aligned peak DA transient (“Dipper Peak”) for saline and J60 groups, respectively. Dashed lines represent linear regressions for each group.

DA responses aligned to lever extension (“Lever ON”) by both J60 and control mice varied significantly with the increasing ratios from 25 to 400, but no overall treatment or interaction effects were detected (RM 2-way ANOVA, n = 5-8, Ratio Factor [F (2.614, 28.76) = 3.742, p = 0.0263], Treatment Factor [F (1, 11) = 0.3206, p = 0.5826], Interaction [F (4, 44) = 1.155, p = 0.3435], Fig. 4E). In contrast, the latency to start pressing was not affected by ratio and it also was not affected by treatment (RM 2-way ANOVA, n = 5-8, Ratio Factor [F (1.897, 20.86) = 2.666, p = 0.0954], Treatment Factor [F (1, 11) = 0.07374, p = 0.7910], Interaction [F (4, 44) = 0.1061, p = 0.9798], Fig. 4 F).

We then analyzed DA responses aligned to dipper elevation (“Dipper ON”) (**Fig. 4G-H**). As in the FR experiment, we observed a similar “double peak” structure and, likewise, the dipper-evoked DA transients increased along with the higher ratios, suggesting that DA release also adapts quickly in response to rapidly changing effort requirements. This adaptation is significantly blunted in the J60 mice (RM 2-way ANOVA, n= 5-8, Treatment x Ratio[F(4, 44) = 3.375,p = 0.0171], Ratio [F(1.814, 19.95) = 39.61, p < 0.0001], Treatment [F(1, 11) = 1.004, p = 0.3378], Fig. 4 I).

Striatal DA release aligned to reward availability (“dipper up”) is thought to encode the immediate value of the effort invested (“sunk”) in obtaining that reward, which can be reflected in the vigor the behavioral response of reaching for the reward after completing the trial. Notably, although there appears to be some modulation of the head entry (HE) latency across ratios for both groups, the RM 2-way ANOVA did not show statistical significance between control and J60 mice (n = 5-8, Ratio [F (2.116, 23.27) = 2.371, p = 0.1131], Treatment [F (1, 11) = 0.1504, p = 0.7051], Ratio x Treatment interaction [F (4, 44) = 0.5798, p = 0.6788], Fig. 4J). However, when we analyzed the relationship between the the DA peaks (aligned to “dipper up”) with the HE latencies throughout the entire session (including all ratios), we found that while control mice displayed a significant negative correlation – the higher the DA peak, the shorter the latency to reach for the reward— (Spearman r = -0.3499, p < 0.0001, Fig 4 K), this relationship was absent in J60 mice (Spearman r = 0.03073, p = 0.5557, Fig. 4 L). Together, these data suggest that in A2a/hM4Ddev mice, DA release associated with a reward-availability cue (dipper up) does not adapt as well to an increasing effort requirement and it is decoupled from the corresponding behavioral response.

## Discussion

Here, we used a viral approach to selectively express the inhibitory DREADD receptor, hM4DGi, in the striatal indirect pathway projection neurons of neonatal A2a-Cre mice. We then inhibited these neurons from postnatal days P8-15 and followed up with behavioral assessments of motivation and NAc dopamine neurotransmission later on. We found that developmentally inhibited A2a/hM4D^dev^ mice exhibited reduced open field locomotion, but only in adulthood, suggesting that the long-term effects of the manipulation do not fully emerge until neurocircuits reach maturity. We further found a decrease in motivated behavior in a progressive ratio task that revealed an impairment in performance by A2a/hM4D^dev^ mice in breakpoint, total overall presses, and session duration. The behavioral results obtained originally with CNO, as the hM4D agonist, were replicated with the JHU37160 (J60). Our analyses did not detect differences in press rate and latency to obtain reward from the magazine, suggesting that the lower PR performance does not reflect a general motor impairment in A2a/hM4D^dev^ mice. Rather, our results suggest a higher sensitivity to the rising cost of each reward that translates into a more conservative approach to effort/reward processing compared to controls. Alternatively, this difference in PR performance could be attributed not to the cost per reward, but to the total effort spent (total number of presses) in a session (830.4 ± 80.6 and 1278 ± 126.2 presses, for J60 and control mice, respectively, Fig. 2). However, when A2a/hM4D^dev^ mice were submitted to the FR40 task during the fiberphotometry experiments (where the total possible press output was 1800 presses, 40 presses/reward in 45 trials), both J60 and control groups completed the task, suggesting that total effort spent does not explain the differences in PR performance. The

We then determined the long-term effects of developmental indirect pathway inhibition on adult dopamine release. In the first assessment of FR block series, DA release was measured across three levels of fixed effort (FR10, FR20, and FR40) done on separate days. Lever extension at the start of the trial serves as a cue that it predicts reward in response to lever pressing. As shown before, such a reward-predicting cue induced a large and sharp DA transient [44] [36]. The DA transient diminished in amplitude with each increasing ratio for both J60 and control groups. Conversely, when DA levels were aligned to dipper elevation, which signals reward availability, the relationship between DA and effort reversed and DA release increased as the effort requirement increased. Alternatively, the increase in DA release is related to the delay to obtain reward or the combination of increased delay and increased effort. These data confirm pervious observations that cue-evoked DA in the NAc adapts with increasing effort requirements encoding upcoming and experienced effort (or time) cost [44]. Importantly, the increase in DA was less pronounced after inhibiting the direct pathway during postnatal development.

In the SR task, mice were exposed to increasing effort requirements <u>within</u> the same session. As for the fixed ratio condition DA transients aligned to lever extension decreased with increasing efforts (though only for low efforts) while transients aligned to the reward availability cue (“dipper up”) increased. Again, the adaptation in DA release to the reward availability cue suggests that DA release at dipper up may reflect spent (sunk) effort and/or may serve as a rapidly updating utility signal that allows the animal to adapt to the changing contingency rules for the upcoming trials. Since the expected reward is always the same at dipper presentation this interpretation stands in contrast with evidence that points to reward-cue NAc DA release primarily encoding future expected reward [45, 46] but also demonstrates a role in encoding spent effort or time [44]. Studies that have applied principles of behavioral economics to independently manipulate both reward value and cost while measuring DA release suggest that reward-cue DA release encodes both effort cost and reward value at the same time, thus helping guide effort-based decisions by providing an integrated index of cost-benefit [47, 48].

In the context of the developmental model our results demonstrated that the developmentally inhibited A2a/hM4D^dev^ mice show an effort-dependent blunted DA response to cost. Our analysis also revealed an expected negative correlation between reward-cue DA peak and the latency to enter the food port for control mice. However, this relationship was abolished in A2a/hM4D^dev^ mice in the developmentally inhibited mice. Thus, instead of a general deficit of DA release, in A2a/hM4D^dev^ mice the DA system appear to be less adaptive to rapidly increasing costs, which in turn could render the mice less able to mobilize their behavior compared to controls.

Our analysis showed that A2a/hM4D^dev^ mice do not display a general deficit of DA release, but that DA is released less only after high effort spent. This contrasts with the findings of the original D2-OE^dev^ model of striatal D2R overexpression where DA release in the NAc was generally reduced [28, 42]. A decrease in VTA DA-neuron firing and NMDA receptor subunit expression in mesolimbic DA neurons observed in anesthetized mice and post-mortem tissue pointed to a general intrinsic deficit in mesolimbic DA function [27].

However, Duvarci and colleagues also measured less recruitment of DA neurons in a cognitive working memory task and found that task parameters regulated DA neuron activity in D2-OE^dev^ mice, which suggested an altered extrinsic regulation of DA neurons depending on the behavioral condition [42]. Since we did not analyze DA neuron activity or NMDA receptor expression in A2a/hM4D^dev^ mice, it is unclear whether the changes in DA release measured in the FR and SR schedules involves intrinsic, extrinsic or both mechanisms.

Differences in the regulation of DA release between both models may be due to the much longer manipulation of D2R expression in D2R-OE mice that started shortly before birth and lasted until adulthood [26]. Alternatively, while both D2R and hM4DGi signal via Gi and arrestin differences in the downstream recruitment of GPCR effectors could account for the differences in both models. Moreover, in D2R-OE overexpressing mice D2R function is increased in response to natural DA release patterns while in A2a/hM4D^dev^ mice the designer receptor is activated twice a day by systemic ligand injection.

Postnatal developmental inhibition of direct and indirect pathways (P8-15) has been shown to change excitatory corticostriatal inputs onto spiny projection neurons, indicating that recurrent circuit activity by both striatal projections via the cortex contribute to synaptogenesis [17]. While developmental inhibition of the direct pathway resulted in decreased frequency of excitatory drive onto and reduced dendritic spine density of striatal projection neurons on P15, the opposite was observed for inhibition of the indirect pathway. Moreover, inhibiting corticostriatal projections during the same period recapitulated the synaptic effects of inhibiting striatal projection neurons. This effect lasted up to 10 days after the manipulation (P25), suggesting that the circuit-level restructuring extends beyond the manipulation. Our results provide evidence that inhibiting the indirect pathway during the same developmental window is sufficient to decrease motivated behavior and open field locomotion well past the manipulation period, 75 days later in adult mice. Surprisingly, open field activity was not changed at P21 suggesting that increased cortico-striatal input at P21 may not affect locomotion.

However, even if cortico-striatal input strengths is enhanced after developmental inhibition of the indirect pathway it is unclear how this may affect the downstream DA system since both direct and indirect pathways are equally affected. One possibility is that the postnatal indirect pathway inhibition leads is not affecting the DA system via recurrent cortical circuitry but by more direct disinhibition of dopamine neurons [49]. This early disinhibition may lead to an adaptive long-lasting decrease in DA neuron excitability thereby reducing the ability of external inputs to activate DA neuron activity. Alternatively, decreased DA release in A2a/hM4D^dev^ mice may involve local regulation of DA release within the NAc which has been shown to modulate motivated behavior [50]. Future studies should be able to address these mechanistic hypotheses.

The first 4 postnatal weeks in rodents constitute a key period of active cellular maturation in the striatum regulated by gene transcriptional networks that determine cell fate and DA receptor molecular identities [51]. In vitro studies have shown that activation of D1R and D2R receptors modulate neurite growth and spine densities during early postnatal development [52, 53] . A programmed increase in DA transmission during early postnatal development has been shown to precede and modulate electrophysiological maturation of SPNs and lack of striatal DA during development leads to immature, hyperexcitable D1-SPNs, which can be prevented by developmental replacement of L-DOPA [16].How increased Gi- or arresting signaling during P8-15 delays or impacts the maturation of striatal projections neurons remains to be addressed in the future.

PET imaging has shown that patients at risk for schizophrenia show enhanced dopamine release capacity during adolescence [11] and extensive DA function during adolescence has been discussed in the etiology of schizophrenia [54]. However, it is unclear how early abnormalities in the DA system arise in schizophrenia since both schizophrenia risk and changes in the DA system are difficult to assess in children. Human genetic studies identified a combination of moderate-risk polymorphisms in DA-related genes and disruptive early life events that contribute to the risk of schizophrenia [55, 56] and mutations in *Drd2a* gene have been associated to cortical thinning and anhedonia [57]. However, causal link between the genetic risk, neural development and disease pathology remains to be shown. Here, we provide evidence that early postnatal inhibition of indirect pathway neurons, as would be expected from enhanced activation of D2Rs by DA, leads to long term deficit in motivated behavior, a negative symptom of schizophrenia, opening the possibility that early changes in the DA system may have long-lasting pathological consequences.

## Acknowledgements

We thank Vivian Zhou for assistance with immunohistochemistry.

## Author Contributions

P.O., P.B. and C.K. conceived the experiments, P.O, M.S. and S.M. performed all experiments.

A.H. developed MATLAB data analysis scripts. R.R. developed protocol and analyzed striatal cell counting. P.O. and C.K. wrote the manuscript with inputs from P.B.

