## Supplementary Figures for "Inhibition of striatal indirect pathway during second postnatal week leads to long lasting deficits in motivated behavior"

**Supplemental Figure S1:**

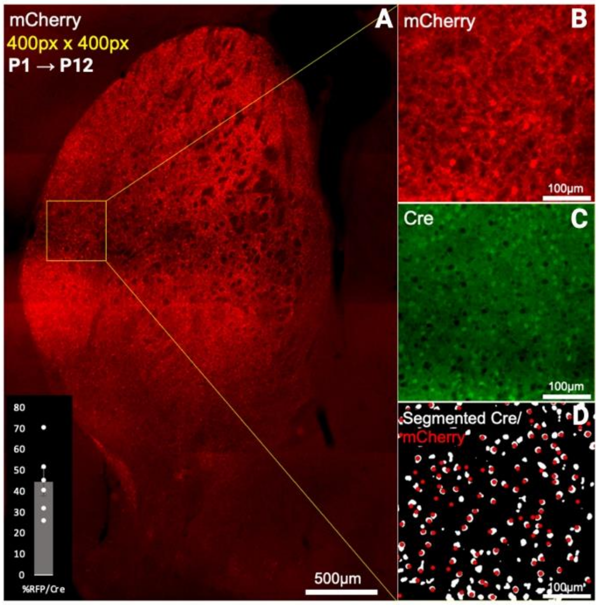


**Figure S1. Viral expression of hM4D-mCherry in A2a-Cre mice. A)** P1 injection of AAV-DIO-hM4D-mCherry in A2A-Cre mouse enables dense labeling throughout striatum, captured here at P12. Yellow square indicates 400 pixel x 400 pixel (410.67µm x 410.67µm) sample used for colocalization analysis. Scale bar 500µm. **B-D)**: Higher magnification images of the region encapsulated in yellow. Shown are single channel examples of immunostained mCherry (panel B) and Cre (panel C). **D)** Detected mCherry cells (red) superimposed on Cre-positive cells segmented (white) by machine learning (see methods). Scale bars 100µm. **Left inset:** Quantification of AAV expressivity, as proportion of RFP and Cre co-labeled cells over total Cre labelled within each 400px x 400px region, plotted as Mean ± SEM (Mean = 44.48876, SEM=6.462696, 6 sections from N=2).

**Supplemental Figure S2:**

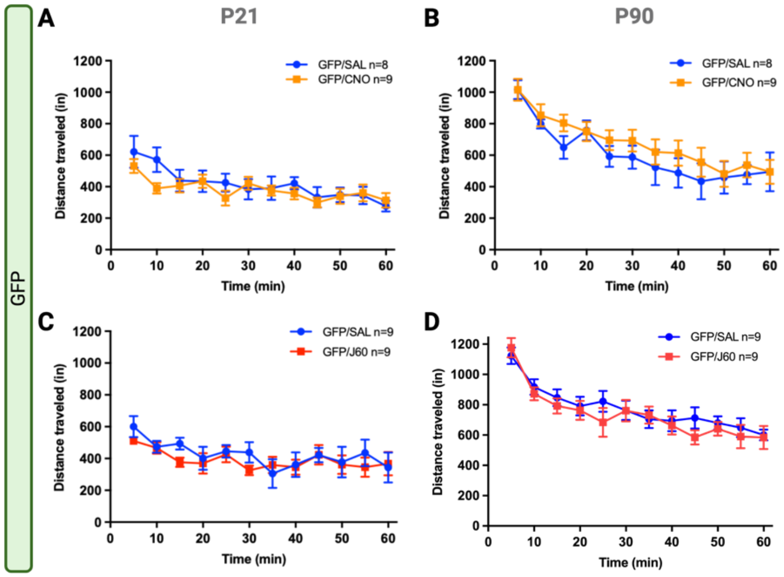


**Figure S2: Open Field test in A2a/GFP$^{dev}$ mice developmentally treated with CNO or J60 (P8-P15) across two different ages. A and B)** Distance traveled over 60 minutes in the OF arena by A2a/GFPdev mice treated with CNO at P21 and P90, respectively, showing no effect of treatment or interaction (RM 2-way ANOVA). **C and D)** Distance traveled over 60 minutes in the OF arena by A2a/GFPdev mice treated with J60 at P21 and P90, respectively, showing no effect of treatment or interaction (RM 2-way ANOVA).

**Supplemental Figure S3:**

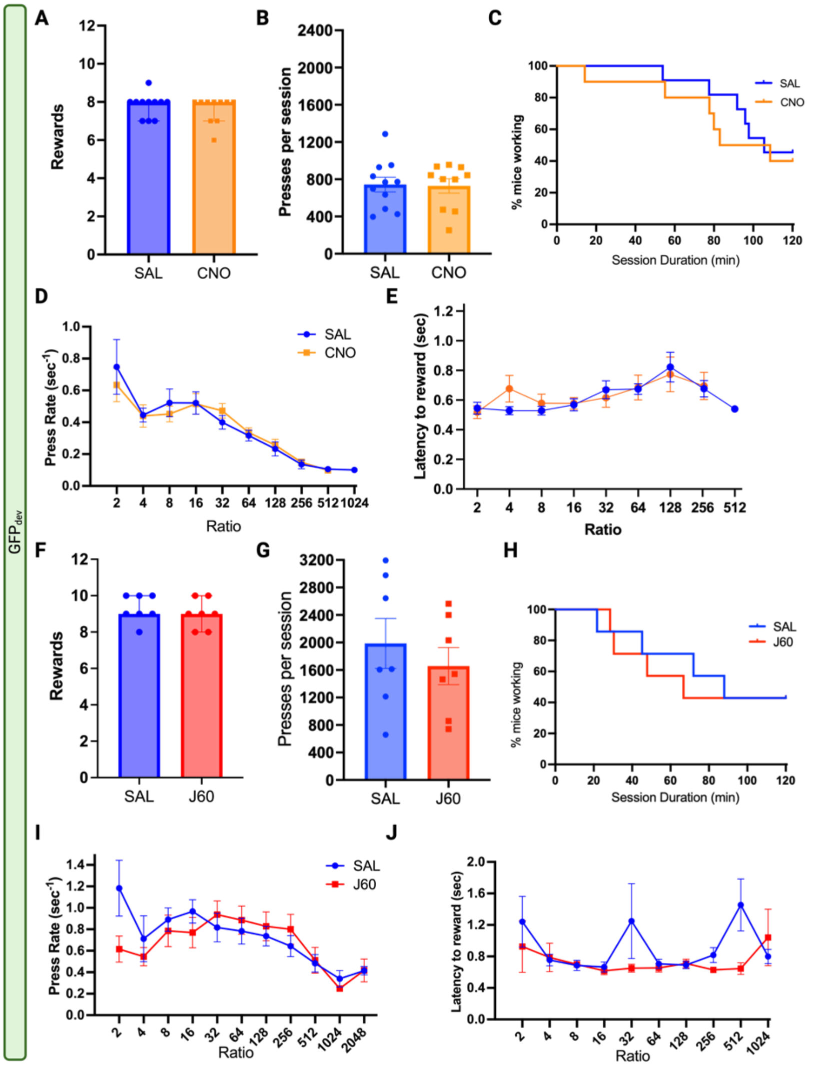


**Figure S3.** ***Progressive Ratio analysis of A2a/GFP$^{dev}$ mice developmentally treated with CNO or J60 from P8 to P15.*** **A-E)** PR results for CNO-treated A2a/hM4D$^{dev}$ mice, n=10-11. **A)** Session breakpoint (earned rewards) for saline- and **CNO**-treated mice. Data expressed as median ± interquartile range, 2-tailed unpaired Mann-Whitney test, p=0.5763. **B)** Summary of total presses per session for saline- and CNO-treated mice, expressed as mean ± SEM, 2-tailed unpaired t-test, p=0.8989. **C)** Survival analysis of percentage (%) of mice engaged in task as a function of session duration. Mantel-Cox Log-rank test, p=0.6681. **D)** Mean (± SEM) press rate (sec$^{-1}$) as a function of ratio, Mixed Effects [(Ratio effect, p<0.0001), (Treatment effect, p=0.8916)]. **E)** Mean (± SEM) latency to reward as a function of ratio, Mixed Effects [(Ratio effect, p=0.0004), (Treatment effect, p=0.8851)]. **F-J)** PR results for **J60**-treated A2a/hM4D$^{dev}$ mice, n=7. **F)** Session breakpoints (earned rewards) for saline- and CNO-treated mice. Data expressed as median ± interquartile range, 2-tailed unpaired Mann-Whitney test, p=0.6999. **G)** Summary of total presses per session for saline- and CNO-treated mice, expressed as mean ± SEM, 2-tailed unpaired t-test, p =0.4797. **H)** Survival analysis of % of mice engaged in task as a function of session duration. Mantel-Cox Log-rank test, p=0.8537. **I)** Mean (± SEM) press rate ($sec^{-1}$) as a function of ratio, Mixed Effects [(Ratio effect, p=0.0024), (Treatment effect, p=0.0646)]; . **J)** Mean (± SEM) latency to reward as a function of ratio, Mixed Effects [(Ratio effect, p=0.1719); (Treatment effect, p=0.2368)].
